# Revealing complex relationships between sex, individuals’ phenotype variation and social signalling in foraging strategies

**DOI:** 10.1101/2024.11.19.624398

**Authors:** Ana V. Leitão, Caterina Funghi, Paulo G. Mota

## Abstract

In social living animals, individuals typically use two main strategies to find food: either by exploiting social information (scrounger) or relying on personal knowledge (producer). These tactics are often linked to different life-history strategies. Access to foraging patches in hierarchical social groups may constrain the use of the producer-scrounger strategies, selecting for colour ornaments to act as status badges. In sexual dimorphic species, males and females may exhibit distinct life-history strategies, with females often lacking a clear badge-of-status and occupying subordinate roles. Here we address the question of how differences in social signalling and personality influence foraging behaviour and the individual’s decision to play producer or scrounger, using males and females European serins, *Serinus serinus.* Specifically, we analysed how individual traits such as sex, plumage colouration, and boldness, relate with foraging strategies in both solitary and social contexts. In solitary contexts, colourful individuals, regardless of sex, were faster to find food, as were bolder males and shyer females. In social contexts, less colourful males and more colourful females adopted the producer strategy, while males’ foraging ability was influenced by their companion’s boldness. Our results reveal that personality and colour differences are linked to social foraging tactics in serins in a sex-dependent manner, highlighting the adaptive value of individual traits in both sexes, and their potential implications for social evolution.

## 1. Introduction

Environmental challenges significantly shape the adaptive strategies of organisms, and understanding the adaptive value of individuals’ alternative strategies offers insight into the potential and limits of evolutionary responses [1, 2]. These adaptive responses manifest as patterns of morphological, physiological, and behavioural traits that co-vary within a life-history, i.e. “Pace-of-Life Syndrome” [3]. Individual differences in foraging behaviour, which are closely linked to fitness and survival, provide an important context for identifying these alternative strategies that individuals use to cope with environmental challenges.

Studies have increasingly examined how variation in resource-use across landscapes is associated with individual traits [4–7]. For instance, personality traits like boldness, neophilia, activity, and exploration may influence the likelihood of encountering new environmental cues, potentially shaping learning opportunities and foraging behaviours [8, 9]. In group-living animals, these individual differences can also inform foraging strategies, where “scroungers” rely on social information and join conspecifics, while “producers” use personal or acquired information to locate food independently [10, 11]. The tendency to adopt one strategy over another can correlate with traits such as sex [12, 13], condition [14], or personality [15]. For example, in Transvolcanic Jays (*Aphelocoma ultramarine*) more dominant individuals scrounge more [16], whereas male yellow-bellied marmots (*Marmota flaviventer)* tend to adopt a producer strategy [17].

Despite increased interest in the relationship between individual traits and social foraging strategies, findings remain limited and often produce contrasting findings. For example, in rooks (*Corvus frugilegus*), bolder individuals scrounge more frequently than shy ones [18], and in barnacle geese (*Branta leucopsis*), shy individuals scrounge more than bold ones [19]. This suggests that directionality of the relationships between traits such as personality, and foraging tactics may be context specific. However, in species with socially hierarchical access to resources, the variation in producer-scrounger tactics may be limited, as dominance hierarchies regulate access to food patches [20]. In such cases, traits like colouration can serve as dominance signals or badge of status, mediating intraspecific competition over food resources while minimising the costs of direct aggression [21].

The European serins (*Serinus serinus)*, is a small, sexually dichromatic finch that forages in groups and flocks outside the breeding season. Both sexes display a carotenoid-based yellow colouration derived from dietary pigments, though only males exhibit this colouration on the crown. Male plumage colouration is a sexually selected trait preferred by females [22, 23], whereas in females, this trait does not appear to influence mate choice [24]. Groups of male serins stablish steep hierarchies regulated by the chromaticity of their yellow crown [25], while female-only groups also exhibit hierarchical structure, though colouration does not seem to affect rank among females [24]. Additionally, studies show that food access within serin flocks generally follows a sequence, with dominant males feeding first, followed by subordinate males, females, and juveniles [26, 27]. However, no research has yet explored producer-scrounger tactics in this species.

In this study, we used a lab-controlled foraging experiment to investigate how individual traits - specifically sex, colouration, and personality – influence foraging strategies adopted in different social contexts. We first assessed individual variation in colouration and boldness-shyness, along with foraging performance across repeated solitary conditions. Given that serins display a diet-dependent, carotenoid-based status badge, we hypothesised that more colourful and bolder individuals would perform better in foraging, regardless the sex. Carotenoid-based colouration, acquired through diet, is often related with foraging ability [28] and habitat availability [29, 30]. We then assessed foraging performance across repeated social conditions, with a same-sex companion that differed plumage colouration. This approach allowed us to examine the social signalling role of colouration outside of mating contexts, assess the consistency of the producer-scrounger strategies, and explore potential sex differences in signal use within social settings. We hypothesised that if colouration serves as an honest signal, it would predict foraging performance across both solitary and social settings. Further, we expected that less colourful males and females would adopt a producer strategy more frequently, reflecting the steep social hierarchy within this species. We further anticipated that the producer strategy would show higher repeatability than the scrounger strategy and that males would generally forage faster than females due to a greater need to maximise carotenoid intake [30], despite females requiring carotenoids for egg yolk deposition.

## 2. Material and methods

### (a) Housing and Morphological Measurements

In November 2013 we captured 34 adults serins (17 females and 17 males) with mist-nets at a site near Coimbra, Portugal (40110250 N 8330350 W). The birds were then transported to the aviary of the Laboratory of Ethology at the University of Coimbra. The aviary had controlled natural ventilation, an ambient temperature of 20 ± 2 °C, and natural lighting supplemented by artificial lights on a 12:12h light–dark cycle.

Individuals were housed in same-sex groups of 4 or 5 individuals in large bird cages (118 x 50 x 50 cm), with ad libitum access to commercial seed mixture (European Finches Prestige, Versele-Laga, Deinze, Belgium), tap water, and mixed grit supplemented with crushed oyster shell. Each bird was banded with numbered black plastic rings (A. C. Hughes, Hampton Hill, U.K.), which were later replaced by plastic colour rings prior to the experiments to allow visual identification during video analyses. We measured morphological parameters, including tarsus (±0.01 mm) and body mass (±0.5 g) one month before the experiments. Body condition was estimated by the scaled body mass index following Peig & Green [31].

### (b) Spectral Analysis

Before the experiments, on the 28th and 29th of May 2014, we measured plumage reflectance in all males and females across 5 patches: crown, throat, breast, belly and uropygial area. For each patch three readings were taken, which were then averaged. Reflectance measurements were made in the avian vision perception range (320 - 700 nm), using an Ocean Optics USB4000 spectrophotometer paired with a MikropackMini-DT-2-GS light source (Ocean Optics, Dunedin, FL, U.S.A.), with a an optical fibre reflectance probe (Ocean Optics R400-7 UV/VIS), held vertically, positioned 3 mm from the feather surface using a standardised holder to ensure consistent distance and exclude all ambient light (sampling area: 28 mm²). Measurements were calibrated relative to a white (Ocean Optics, WS-1-SS) and dark standards.

Reflectance spectra was summarised using psychophysical models of avian vision [32, 33]. We calculated cone quantum catches for each of the four cone types present in the avian retina, sensitive to very short (VS), short (S), medium (M) and long (L) wavelengths, integrating cone sensitivity, irradiance and plumage reflectance. We assumed that *S. serinus* has the same visual system as its close relative *S. canaria*, which is known to have the UVS visual system [34]. We considered the cone sensitivities of another ultraviolet sensitive (UVS), the blue tit, *Cyanistes caeruleus* [35]. As a measure of irradiance, we used the spectrum of standard daylight, D65 [33].

Plumage colouration was quantified using the short-wavelength-sensitive (SWS) ratio, representing a chromatic index of plumage reflectance [36], similar to Leitão et al. [25]. Visual models were calculated in R [37], using the *pavo* package [38].

To create a composite measure of plumage colouration, we conducted a Principal Component Analysis (PCA) on the SWS values from the throat, breast, belly, and uropygial areas. The first Principal Component (hereafter: ‘colouration’) explained 72.4% of the total variation and, after positive transformation, this component showed moderate positive correlations with belly (0.51), throat (0.52), breast (0.55), and uropygial (0.4) patches.

### (c) Personality tests

We conducted three behavioural tests designed to measure different personality traits in birds: 1) Tonic immobility to assess fear, 2) Mirror test to assess sociability, 3) New object test to assess boldness. All tests were conducted between 0900 and 1300h over two rounds, each including 32 tests (17 females and 17 males), separated by approximately 6 weeks. Due to a video-recording issue, two tests from both the new object and mirror tests were excluded, resulting in a total of 32 tests for each of these.

The tonic immobility test measured the latency for a bird to pass from a stress-induced motor inhibition to a mobile state [39]. Tests were conducted from 19 to 24 March (1st round) and on 4 May (2nd round). Each bird was removed from its home cage and placed in a darkened cover for one minute. Afterward, the bird was placed on its back on a flat wooden surface (a 15.5 × 15.5 cm), elevated 87 cm from the floor, in the centre of an empty room (1.36 × 1.06 × 2.27 m), lying sideways to an observer stationed 40 cm away. The observer recorded the time until the bird returned to an upright state or flew away (latency to fly), with a maximum of 60 seconds for each trial. Birds that did not fly were given a maximum score of 60 seconds.

To assess sociability, we used a mirror test, following previous studies where serins did not display aggressive reactions to mirror reflections. Tests were conducted from 26 to 29 March (1st round) and from 6 to 8 May (2nd round). Each test took place in a modified bird cage (54.3 x 27.5 x 37 cm) with four perches and a mirror (27.5 x 37 cm) on the side, initially covered by a removable cardboard. The test was filmed for 10 min, where the first five minutes the mirror was covered, followed by 5 minutes with the mirror uncovered. Upon exposing the mirror, we recorded the bird’s position and duration of proximity to the mirror, calculating both the proportion of time spent close to the mirror and a position index that weighted time spent in closer versus farther positions.

To measure boldness, we conducted a new object test, similar to other studies [40, 41], that focused on phobia and exploratory behaviour. Tests were conducted from 31 March to 2 April (1st round) and from 10 to 15 May (2nd round), in the individual’s home cage, with 4 perches of regular intervals (Figure 13). Food was removed 1 to 2 hours prior to testing to motivate exploration. For each test, all cage mates were removed, except for the focal individual. We placed a temporary partition within the cage, then introduced an unfamiliar object - a yellow clothes peg - on the perch furthest from the bird, with a food dish placed adjacent. Upon removal of the partition, the bird’s activity was recorded for 10 minutes (600 seconds). To control for potential side biases, the object and feeder position were alternated across trials. We recorded latency to approach the object, and measured movement between perches, generating a position index that emphasised positions closer to the object. Birds that did not approach were given a maximum latency score of 600 seconds. Despite the home-cage setting, we classified “activity between perches” as exploratory due to the novel object, altered food conditions, and absence of cage mates.

We calculated repeatability between rounds (see statistical methods), and only the new object test showed repeatability (Table 1). Therefore, only data from the new object were included in subsequent models.

**Table 1.** Summary of the adjusted repeatability (R) analysis of individuals’ behavioural and foraging tests. The confidence interval (CI) at 95% is provided. Behavioural assay, variable tested, fixed terms and sample size (N) are specified. The values in bold represent the repeatability considered significant, with CI different from zeros (higher than 0.1).

| Behavioural Assay | Variable | Fixed terms | R | CI 95% | N |
| --- | --- | --- | --- | --- | --- |
| Tonic immobility | Latency to fly (log) | Sex | 0 | [0,0.23] | 34 |
| Novel object | Position Index | Sex | 0.16 | [0,0.50] | 32 |
| Novel object | **Latency to new object** | Sex | **0.4** | **[0.1,0.66]** | 32 |
| Novel object | **Activity** | Sex | **0.49** | **[0.21,0.73]** | 32 |
| Mirror | Position Index | Sex | 0 | [0,0.36] | 32 |
| Mirror | Activity | Sex | 0 | [0,0.34] | 32 |
| Mirror | Duration looking to mirror | Sex | 0 | **[0,0.37]** | 32 |
| Mirror | Time close to the mirror (proportion) | Sex | 0 | [0,0.05] | 32 |
| Solitary Foraging | **Latency to eat the first seed (log)** | Sex | **0.44** | **[0.12,0.69]** | 34 |
| Solitary and social foraging | **Latency to eat the first seed (log)** | Sex, PairID | **0.45** | **[0.35,0.88]** | 34 |
| Social Foraging | Latency to eat the first seed (log) | Sex, PairID | 0 | [0.55,0.99] | 34 |
| Social Foraging | **Producer strategy** | Sex, PairID | **0.28** | **[0.72,0.99]** | 34 |

### (d) Foraging Experiments

The foraging experiments were conducted from June 2 to June 29, 2014, between 0930 and 1830, initially with 17 males and 17 females. However, due to temporary illness affecting one male and one female in the later phases, certain tests were excluded, leading to variable sample sizes, which are reported throughout. The foraging experiments were conducted from June 2 to June 29, 2014, between 0930 and 1830, initially involving 17 males and 17 females. However, due to illness in some birds, certain tests were excluded, resulting in variable sample sizes.

To standardise foraging motivation, in each trial individuals were food-deprived for 2h prior to testing in their home cages. Individuals were then transferred to a test room (136 x 112 cm and 230 cm high), illuminated by fluorescent lamps (Philips TLD 36 W/950) with the lights off, with tests beginning immediately after the lights were turned on. The foraging setup consisted of a table (48.5 x 48.5 cm) with 16 evenly spaced holes (1.5 cm diameter, 0.8 cm deep), separated by 8.5 cm (Figure 1). The experiment comprised five sequential trials, each lasting approximately one week. Birds were tested in the same order throughout the stages to maintain consistent timing across trials. In the first trial 1, all 16 holes were filled with 10 seeds each to help the birds to acclimate to the setup. In subsequent trials 2-5, only 4 holes were filled, each containing 20 seeds. The selection of filled holes was balanced and rotated systematically to avoid repetition. We used *brassica napus* seeds, which are part of the serin’s natural diet and known to be a preferred choice in the lab. These seeds contain Lutein and B-caroten [42].

**Figure 1.**
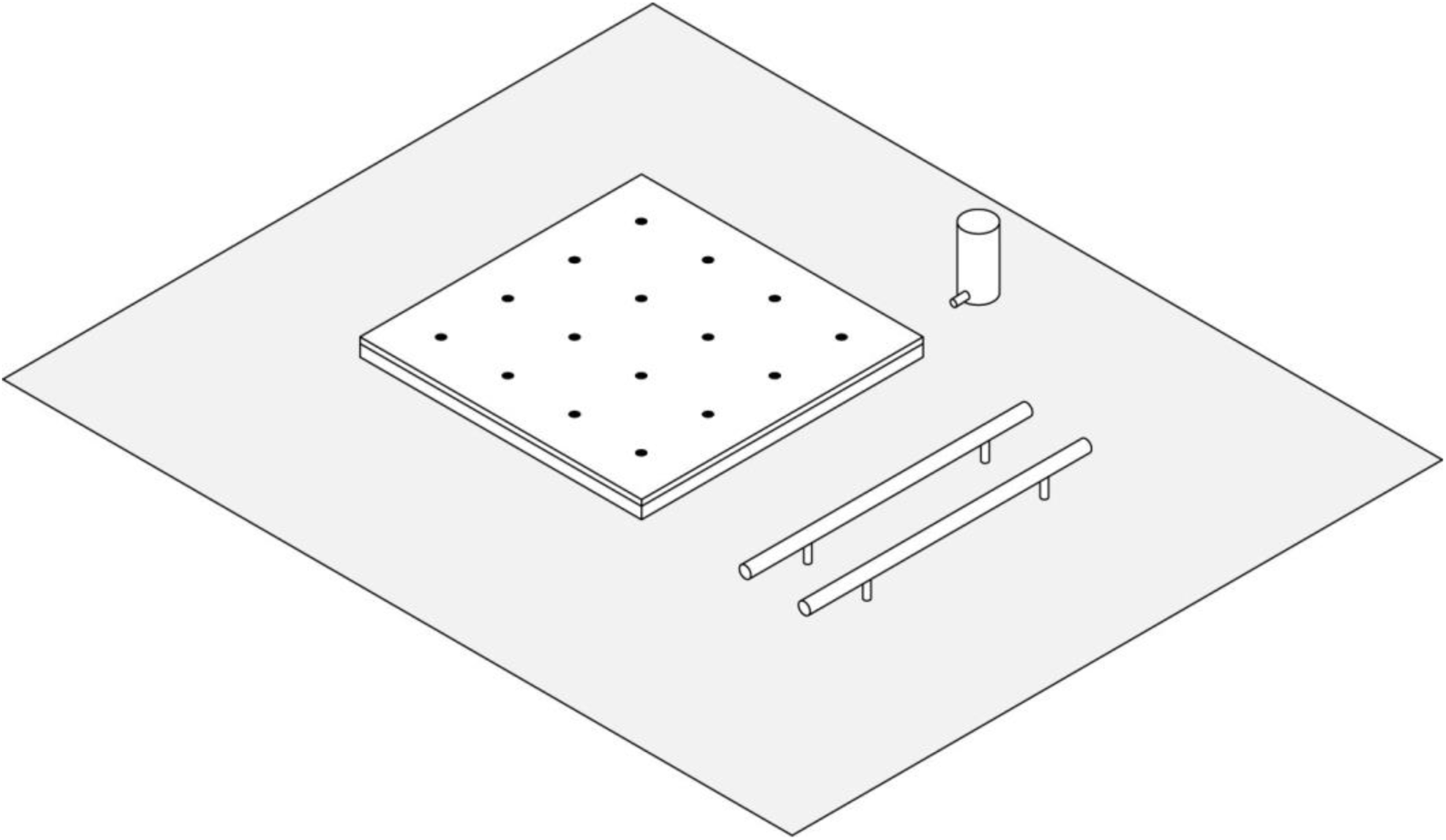
The apparatus for the foraging experiment: the room test had a perch, access to water and a foraging table (48.5 x 48.5 cm) with 16 holes (0.8 dept, 1.5 diameter). The holes were spaced 8.5 cm between them.

Trials alternated between two conditions: solitary (trials 2 and 4) and social (trials 3 and 5). In the social trials, birds were paired by selecting one individual from each colour category (more and less colourful) to allow comparisons based on colour-based social dynamics.

Each trial lasted 1h and was recorded using a Sony SSC-DC378P video camera. Behavioural data was analysed using Observer XT 10.0 software (Noldus Information Technology, Wageningen, Netherlands). We measured latency to find the first hole with seeds as an indicator of foraging performance. Furthermore, we scored the number pecking in the holes with seeds (eating), the number of pecking in the empty holes (searching). In social contexts we also scored the number of pecking holes when an individual was already present (joining). We used these behaviours to calculate the percentage of ‘producer strategy’ following Katsnelson et al. [43], as the proportion of active searching and eating events relative to the sum of searching, eating and joining events.

### (e) Statistical Analysis

#### Repeatability of personality and foraging behaviour

To access the consistency of individual differences in personality (using novel object, mirror and tonic immobility assays) and foraging performance (in both solitary and social context), we ran an adjusted repeatability analysis for each of the variable processed from the videos (two trials for each behavioural or foraging tests). We used (G)LMMs, assigning the proper error structure based on the datatype (i.e. Gaussian, binomial, proportion, or Poisson) with the ‘rptR’ package in R [44]. Each repeatability estimate was assessed with 1000 permutation to determine the likelihood of the 95% confidence interval to be significantly different from zero. To account for potential sex differences, we adjusted repeatability estimates for personality and solitary foraging metrics by sex. Social foraging estimates were adjusted for both sex and pair ID to control for non-independence of the data. Only behaviours that were repeatable were included in further analyses (Table 1). Of all personality tests, only the novel object assay was found to have consistent individual differences across trials (R >0.4, N = 32, Table 1). We summarised the variables of the novel object (latency to approach the food and the number of movements), into a single ‘new object score’ through a Principal Component Analysis, using the mean values across trials. The first principal component (hereafter: ‘boldness’) explained 54% of the total variation, correlated positively with activity (0.7) and negatively with latency (-0.7), indicating that high values of the score reflect bolder individuals.

#### Foraging performance across solitary and social contexts

We assessed whether plumage coloration and boldness influenced individual foraging performance, measured as the latency to find the first hole containing seeds, comparing this performance across two distinct contexts: solitary and social. Using data from the first trials in both solitary (trial 2) and social (trial 4) contexts, we tested whether the latency to find the first hole with seeds was influenced by context (solitary vs social), plumage colouration, boldness score, and sex. We built a Linear Mixed Model (LMM) with Gaussian error structure, with latency to consume the first seed (log-transformed) as the dependent variable. Fixed effects included individual colouration, boldness score, and their interactions with both sex and context as fixed terms. Bird ID and Pair ID were included as random effect to account for individual and pair-level repetition.

#### Social foraging performance

To investigate the role of plumage colouration and personality in social foraging contexts, we analysed the social foraging trials, focusing on the producer strategy (calculated as the number of eating and searching events relative to the total number of eating and searching events). A GLMM with binomial error structure was used, with producer strategy as the dependent variable. Fixed effects included individual boldness, score the individual colour category (i.e. more or less colourful than the foraging companion), the foraging partner’s boldness score and sex interaction with these variables, and the difference in scaled body mass index between focal individuals and their foraging partners to control for body size differences. Pair ID and Bird ID were fitted as random terms to account for individual and pair-level repetition.

In all models, we employed a stepwise reduction process sequentially removing the least significant terms beginning with interactions, as determined by likelihood ratio test between models. The “lme4” package [45] was used for model building, and post-hoc interactions between categorical variables were tested using the “phia” package [46]. All analyses were conducted in R [37].

### 3. Results

On average, males approached the novel object faster than females, with a latency of 346.28 ± 241.53s for males compared to 428.36 ± 216.71s for females (average for both rounds). In solitary foraging trials, males also outperformed females in finding the first seed (average latency to find the first seed: males = 1307.94 ± 1213.67s, females = 1451.48 ± 1124.93s). In social foraging trails, with a foraging companion, the difference between sexes disappeared (males = 545.29 ± 870.62s, females = 532.90 ± 893.98s). Latency to eat the first seed was found repeatable across solitary foraging trials (R = 0.44), and solitary and social foraging contexts (R = 0.45), but not across social foraging trials (Table 1). The producer strategy was found repeatable across social trials (R = 0.28, Table 1).

When examining individual foraging performance across both solitary and social contexts, we found that more colourful individuals were faster at finding food when foraging alone compared to less colourful ones, while in social context, this effect was weaker, with more colourful individuals taking slightly longer to find food (interaction between colouration and foraging context significantly influenced latency: Figure 2a, Table 2, N_obs_=62, N_ID_=32). Furthermore, boldness influenced foraging performance differently in males and females regardless of context: bolder males were faster finding food than shy ones, while bolder females were the slowest to find the first seed (Figure 2b, Table 2, N_obs_=62, N_ID_=32).

**Figure 2.**
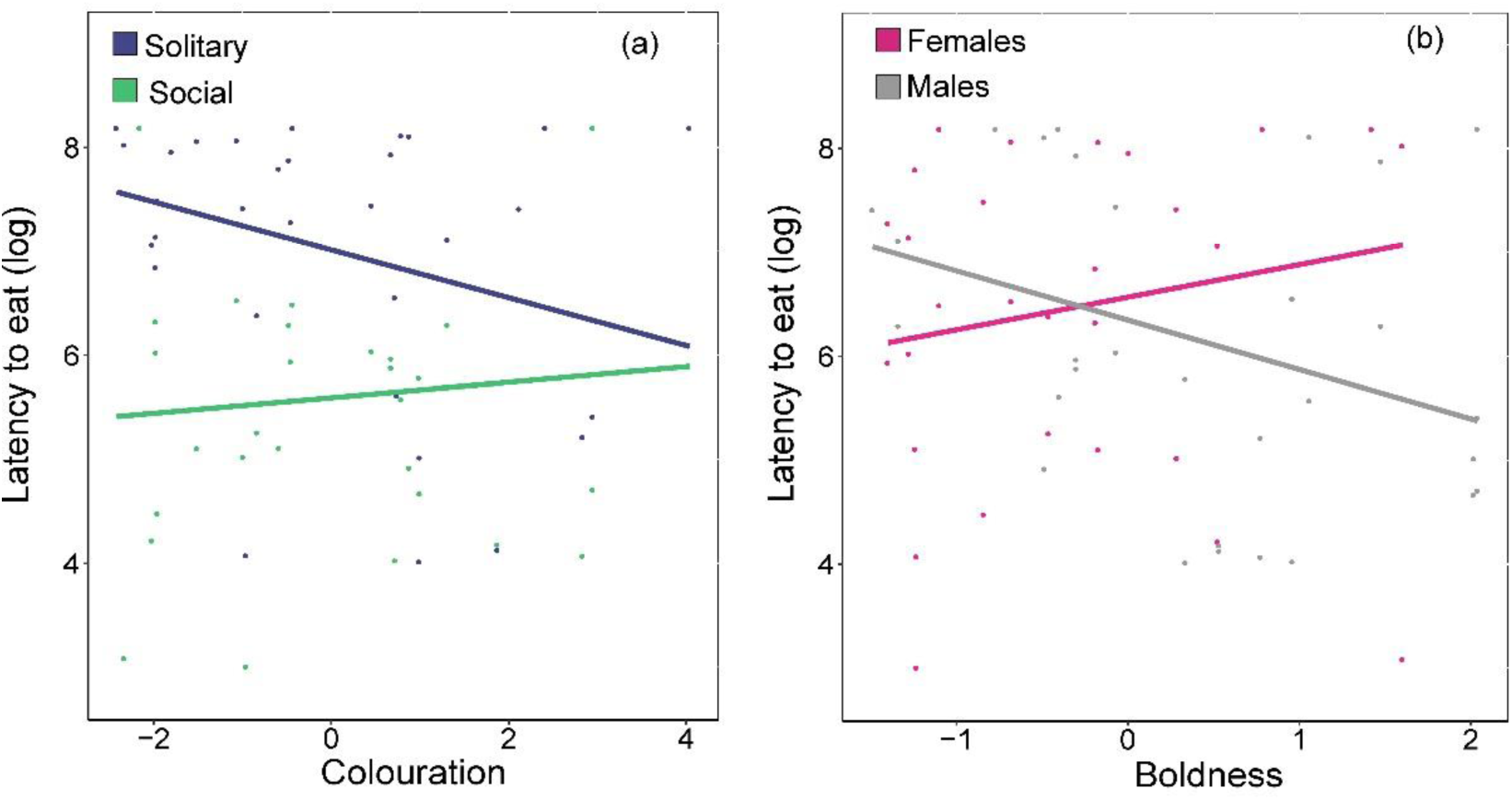
Graphical representation of the model with individuals foraging behaviours in solitary and social contexts. Relationship between a) latency to eat the first seed (log-transformed) and colouration, b) latency to eat the first seed (log-transformed) and boldness. Significant interactions are showed with blue and green (lines and dots) which represent solitary and social contexts respectively, while grey and pink (lines and dots) represent males and females, respectively. All the lines represent model predictions.

**Table 2.** Summary of the (G)LMMs. Significant p-values (P) are highlighted in bold. Response variables, random terms sample size (N) and variance (Var) are specified for each model. Value of fixed effects (Estimate) and standard errors (S.E.) are estimates for the variables in minimal adequate model; values in brackets represent coefficients and S.E. from the model before the term was dropped. The coefficients of the test (X^2^) and the degrees of freedom (d.f.) of non-significant categorical variables are not specified.

| Context | Response variable | Random term | N | Var | Fixed effect | Estimate | S.E. | $X^2$ (d.f.) | P |
| --- | --- | --- | --- | --- | --- | --- | --- | --- | --- |
| Solitary vs Social | Latency to eat (log) | Bird ID | 32 | 0.57 | (Intercept) | 5.95 | 0.45 |  |  |
|  |  | (Residuals) |  | 0.97 | **Context\*Colouration** |  |  | 4.99(1) | **0.03** |
|  |  |  |  |  | **Sex<sup>a</sup>\*Boldness** |  |  | 4.02(1) | **0.05** |
|  |  |  |  |  | Colouration | 0.23 | 0.23 |  |  |
|  |  |  |  |  | Sex<sup>a</sup> | -0.42 | 0.77 |  |  |
|  |  |  |  |  | Boldness | 0.34 | 0.29 |  |  |
|  |  |  |  |  | Context<sup>b</sup> | 1.37 | 0.25 |  |  |
|  |  |  |  |  | Context<sup>b</sup>*Colouration | -0.35 | 0.15 |  |  |
|  |  |  |  |  | Sex<sup>a</sup>*Boldness | -0.79 | 0.38 |  |  |
|  |  |  |  |  | Context<sup>b</sup>*Boldness |  |  | 0.28(1) | 0.59 |
|  |  |  |  |  | Sex<sup>a</sup>*Context |  |  | 0.28(1) | 0.59 |
|  |  |  |  |  | Sex<sup>a</sup>*Colouration |  |  | 0.02(1) | 0.88 |
| Social | Producer strategy | Bird ID | 30 | 3.43 | (Intercept) | 1.53 | 1.01 |  |  |
|  |  | Pair ID | 27 | 4.19 | **Colour category\*Sex<sup>a</sup>** |  |  | 10.65(1) | **0.001** |
|  |  |  |  |  | **Boldness companion\*Sex<sup>a</sup>** |  |  | 10.71(1) | **0.001** |
|  |  |  |  |  | Colour category<sup>c</sup> | 2.45 | 1.04 |  |  |
|  |  |  |  |  | Boldness Partner | -1.44 | 0.54 |  |  |
|  |  |  |  |  | Sex<sup>a</sup> | 1.33 | 1.37 |  |  |
|  |  |  |  |  | Colour category<sup>c</sup>*Sex<sup>a</sup> | -5.06 | 1.46 |  |  |
|  |  |  |  |  | Boldness Partner*Sex<sup>a</sup> | 2.19 | 0.63 |  |  |
|  |  |  |  |  | Boldness | (-0.2) | (0.45) | 0.19(1) | 0.65 |
|  |  |  |  |  | Scaled body mass index difference | (-0.1) | (0.23) | 0.14(1) | 0.71 |
|  |  |  |  |  | Sex<sup>a</sup>*Boldness |  |  | 1.42(1) | 0.23 |

Categorical terms: <sup>a</sup> Sex: the reference is male; <sup>b</sup> Context: the reference is Solitary; <sup>c</sup> Colour Category: the reference is more colourful

Less colourful males adopted the producer strategy, i.e. actively searching and eating the seeds, more frequently than the more colourful ones (males: colour category=0.93, *X*^2^=6.47, P=0.02), while in females the opposite pattern was observed (females: colour category=0.08, *X*^2^=5.54, P=0.04, Table 2, Figure 3a). Additionally, more colourful females had higher producer strategy than more colourful males (more colourful category: females-males= 0.98, *X*^2^=8.47, P=0.007, Figure 3a). Furthermore, the interaction between boldness companion and sex was significant in influencing producer strategy (Table 2): males paired with a bolder companion showed an increased tendency to adopt a producer strategy, a pattern that was not observed in females (Figure 3b, N_obs_=57, N_ID_=30, N_pair_=27).

**Figure 3.**
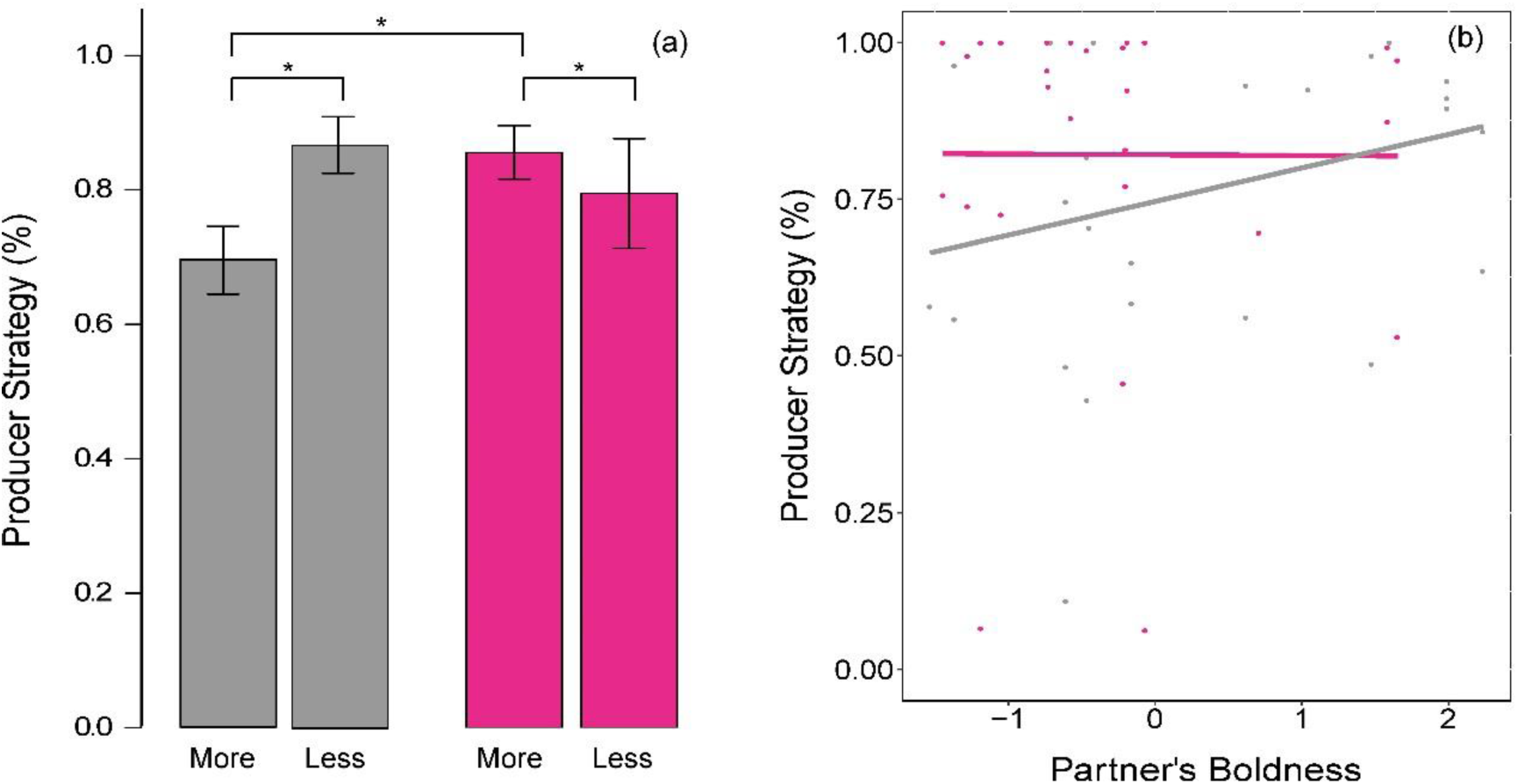
Graphical representation of the model with individuals’ percentage of producer strategy used to forage with a foraging partner. (a) Mean ± S.E. producer strategy used by the more and less colourful males (grey) and females (pink). Significant differences are marked with *. (b) Relationship between producer strategy and the interaction between foraging partner’s boldness and sex. Lines represent model predictions. Grey and pink lines and dots represent males and females, respectively.

## 4. Discussion

Our findings reveal that individual traits such as sex, colouration, and personality significantly influence foraging behaviour and strategies in European serins. We found that carotenoid-based ornamentation reliably signals foraging ability in both sexes, with more colourful individuals performing better under solitary conditions, while less colourful individuals excelled in social conditions. Additionally, bolder males and shyer females showed higher foraging performance, highlighting a sex-specific relationship between personality and foraging success. In social foraging contexts, less colourful males and more colourful females adopted the producer strategy, with male tactics influenced by the boldness of their companions, an effect not observed in females. To our knowledge, this is the first study to experimentally test the effects of variation in colouration and personality traits in foraging performance across both sexes in different social contexts.

### (a) Impact of colouration, sex and boldness on individual foraging performance

Our results indicate that carotenoid-based ornamentation serves as a reliable predictor of foraging ability in solitary contexts for both male and female European serins. This suggests that individuals with greater colouration have better access to high-quality food resources, consistent with findings linking carotenoid-based ornamentation and foraging ability [28, 30, 47, 48]. This also reflect an underlying link between their physiological condition and foraging efficiency. Few studies, however, have examined ornamentation’s role in both sexes, making our research one of the first to do so (but see [49].

The relationship between personality traits and foraging performance is complex and varies across taxa, with boldness often linked to faster or greater resource monopolisation [19, 50, 51]. Our findings highlight the complexity of the relationship between sex, personality traits and foraging performance: bolder males and shyer females were faster at finding the first hole with seeds and had higher foraging performance. This suggests that males and females employ different strategies to optimise foraging based on distinct personality traits. Sex-related differences in personality combined with cognitive abilities, can influence foraging strategies [52, 53], and may reflect sex-specific covariances in selective pressures shaping life-history strategies, physiological adaptations, and behaviours [3, 54].

### (b) Context-Dependence of Foraging Strategy

In species like serins, with steep hierarchical access to food is often regulated by ornamental signals, favouring dominant individuals and enforcing sex differences [26]. Although both male and female colouration reliably indicates solitary foraging ability, social contexts favour different strategies based on colouration and sex. Less colourful individuals generally outperformed more colourful ones in locating food within group settings, with less colourful males and more colourful females more frequently adopting producer roles. This suggests that colouration as status signal primarily benefits males, since with a foraging companion, more coloured males were slowest to find the first seed and tended to scrounge more. By contrast, females did not take advantage of their colouration in social context, although it reliably reflected higher foraging performance in solitary conditions. These results may illustrate possible mechanisms in the evolution of social signalling, wherein ornaments minimise the costs of aggressiveness to food access in social context [21]. Females, characterised by less ornamentation in serins, may face trade-offs between the costs of maintaining ornaments with the ones of reproduction [55] and thus suffering in group foraging scenarios.

Furthermore, our results suggest that the consistent preference for adopting a producer strategy, over scrounging, may be explained by the hierarchical access to food, with scrounger tactics being more tolerated among less colourful males and females. In granivorous species, food sources are typically widespread and abundant, though dominant individuals may still engage in scrounging if it involves aggressive displacement [26, 56].

The interplay between boldness and social interactions significantly influences foraging tactics in group-living animals. Our findings indicate that males paired with bolder companions were more likely to adopt a producer strategy, suggesting that a competitive environment created by bold companions encourages others to leverage their personal knowledge and to adopt a riskier but potentially more rewarding foraging strategy. Importantly, these results highlight that foraging behaviour is not solely determined by individual traits but is also influenced by the characteristics of companions within the social group.

In contrast, female foraging performance was not affected by the companion boldness, suggesting that in social contexts, females may prioritise their own foraging abilities over the influence of others. Furthermore, females may benefit more from minimising predation risks than from the advantages of social foraging.

These findings illustrate the intricate relationship between sex, boldness, and foraging tactics, reflecting a fine balance between cooperative and competitive strategies in group-living animals. Such dynamics can have significant implications for the evolutionary pathways and ecological interactions of these species. Future research should further explore these sex-specific strategies, considering how they affect overall group foraging efficiency, resource allocation, and the potential for social learning.

Overall, our research shows how individual traits influence foraging strategies in European serins (*Serinus serinus*), highlighting how sex and foraging ability mirror different life-history strategies and concur to explain phenotypic variability. Depending on their sex, personality type, and social context, individuals may experience and adopt different foraging strategies. This variation may facilitate adaptation to fluctuating environments, allowing individuals to leverage different strategies based on their position within a social hierarchy and environmental cues. By elucidating these individual differences in foraging behaviour, we can better understand the adaptive value of individual traits, reinforcing the idea that successful foraging is not merely a function of environmental availability but also of individual characteristics.

## Ethics

This work was performed in compliance with the regulations of the Portuguese National Authority for Animal Health (DGAV). All permits for animal capture, transport, maintenance, handling, and experimental procedures were carried out under I.C.N.F. licenses (43, 44 and 45/2013/CAPT) according to Portuguese legislation.

Birds were initially captured and transported to the aviary in small groups in cages (70 x 35 cm x 50 cm). Upon arrival, birds were moved to their designated housing cages, where their welfare was monitored daily to ensure their health and well-being. Additionally, a veterinary surgeon also checked the birds’ general health and their housing conditions. After the experiments, birds were transferred to a large indoor aviary for a 5-day period of flight training, to restore flight performance and activity levels to those typical of wild birds. Individuals were then released at their capture locations, where they rejoined local groups of conspecifics.

## Data accessibility

Behavioural data as well as R scripts will be available at Dryad upon publication.

## Declaration of AI use

AI-assisted technologies were used to help for English language support and text editing (Confluence and ChatGPT). However, all content was carefully reviewed and edited by the authors, and we assume full responsibility for its final form.

## Authors’ contributions

A.V.L.: conceptualisation, data curation, investigation, methodology, project administration, writing—original draft, writing—review and editing; C.F.: investigation, methodology, formal analysis, visualisation, writing—original draft, writing—review and editing; P.G.M.: funding acquisition, conceptualisation, methodology, project administration, supervision, writing— review and editing.

All authors gave final approval for publication and agreed to be held accountable for the work performed therein.

## Conflict of interest declaration

We declare we have no competing interests.

## Funding

This research was funded by the Portuguese Research Council (FCT), through a project PTDC/BIA-BEC/105325/2008 to P.G.M.

## Acknowledgments

We are grateful to Eliana Soukiazes for her assistance with bird captures, maintenance, and colour measurements, as well as to the undergraduate Biology students from the 2014 cohort for their help with video analysis. Special thanks to Eng. António M. Neto for building the foraging board used in this study.

## Notes

### Competing Interest Statement

The authors have declared no competing interest.

